# Male success against con- and heterospecific competitors indicates a positive but modest role for sexual selection as driver of speciation

**DOI:** 10.1101/231852

**Authors:** Jeremy S. Davis, Leonie C. Moyle

## Abstract

For sexual selection within species to drive the evolution of reproductive isolation between lineages, sexually selected and reproductive isolating traits must both share underlying mechanisms and operate in the same direction. While some work has been done to evaluate mechanistic overlap, fewer studies have evaluated whether intraspecific sexually-selected variation is associated with elevated reproductive isolation between species. Here we evaluate this association by assessing the relationship between male reproductive success against conspecifics versus heterospecific males at each of two different mating stages. We find that male precopulatory performance (remating success following a conspecific versus a heterospecific first mating) was not associated between conspecific and heterospecific contexts, but postcopulatory success (sperm competition against conspecific versus heterospecific males) was modestly positively correlated. We discuss two lines of evidence that suggest this modest association is due to incomplete mechanistic overlap between postcopulatory competition in conspecific and heterospecific mating contexts. This study provides an explicit test of a necessary condition for sexual selection to drive speciation, and finds that while sexual selection is not individually sufficient to explain the magnitude of reproductive isolation in this system, it could nonetheless facilitate the evolution of isolation via postcopulatory sperm competition.

## Introduction

Sexual selection is frequently proposed as a powerful driver of speciation (Ritchie 2007), however for this to be the case, two conditions must be met. First, traits that are the target of sexual selection must also be involved in reproductive isolation, so that the two processes share mechanisms and underlying genes in common. Second, sexual selection and reproductive isolation must act in the same direction. In particular, in order to drive speciation, sexual selection must favor trait changes within species that act to amplify reproductive isolation between species. Some empirical studies have generated evidence that sexual selection and species reproductive isolation act via shared traits, mechanisms, and/or genes (Groot et al. 2013, Arbuthnott 2009, Castillo and Moyle 2014, and see below), although whether they are sufficiently mechanistically coupled for sexual selection alone to drive isolation remains equivocal (Panhuis et al. 2001, Ritchie 2007, Bolnick and Kirkpatrick 2012, Safran et al. 2013). Moreover, it is equally unclear that the direction of sexual selection and reproductive isolation will consistently align; theory suggests that they might act at cross purposes under some conditions (Servedio and Burger 2014), but there are few empirical studies that explicitly examine the strength and direction of association between traits that mediate sexual success within and between species.

A range of sexual traits could potentially be involved in both intraspecific reproductive performance and reproductive isolation against heterospecifics. In particular, studies of precopulatory traits (reviewed Ritchie 2007), especially male traits related to courtship, have produced some evidence that male signal traits that are strongly preferred by conspecific females also strongly contribute to interspecies mating isolation (e.g. *Laupala* crickets: Mendelson and Shaw 2005, Shaw and Parsons 2002). Nonetheless, other less examined mating interactions might also play an influential role in both intraspecific mating success and interspecific reproductive barriers. Remating in *Drosophila* offers a context within which to investigate multiple such mating traits with consequences for both conspecific and heterospecific interactions. Among *Drosophila*, remating is a common sexual strategy. In *Drosophila melanogaster*, paternity tests from natural field collected females indicate that most have mated between 2 to 6 times (Imhof et al. 1997, Harshman and Clark 1998), while remating estimates are >80% in laboratory settings (reviewed Singh et. al. 2002). Accordingly, the majority of matings are expected to be rematings, making male performance in remating contexts an important aspect of lifetime fitness. To secure offspring with a female that has previously mated, a second male needs both to successfully court this female and, once mated, to effectively displace the sperm of the first mated male (referred to as ‘offensive’ sperm competition; Boorman and Parker 1976). In the initial (pre-copulatory) interaction, the male must convince the female—via acoustic, visual, and chemical cues—to accept a second mating, despite potentially detrimental effects to her and despite biochemical manipulation by the first male that decreases her receptivity (Parker and Partridge 1998, Sirot et al. 2009). Because remating provides females with a degree of control over paternity of offspring, especially when the first male is deemed suboptimal (Fricke et al. 2006), female remating decisions likely involve an assessment of the quality of the second male. In interactions with conspecifics alone, this assessment can be based on factors such as sperm depletion (when a first mating was not recent; Clark et al. 199, Gromko and Markow 1993) but also on optimizing mating with the highest quality males (‘trading up’ Byrne and Rice 2005). However, it’s unclear whether the same performance characters would be assessed, or assessed to the same degree, by females whose first mating was with a heterospecific male—a mating context that is almost always suboptimal. If females evaluate different qualities when choosing remating partners depending on whether a first male was conspecific or heterospecific, sexual selection on precopulatory remating traits need not act in the same direction as reproductive isolation.

Following a successful remating event, a second male must also outcompete the first male in sperm competition. In *Drosophila*, it is well documented that the second (or ‘P2’) male frequently sires more offspring than the first (‘P1’) male (a phenomenon referred to as ‘P2 precedence’), whether the first mating is conspecific or heterospecific (Price et al. 1999, Price 1997). Nonetheless, P2 success against conspecifics can vary among males, depending upon the factors such as condition and sperm count (Letsinger and Gromko 1985) as well as female genotype (Clark et al. 1999, Bjork et al. 2007). In contrast to intraspecific sperm competition, second mated conspecifics almost always sire the majority of offspring following a first heterospecific mating, a phenomenon known as conspecific sperm precedence (Price 1997). This phenotypic observation suggests that sperm competition between conspecifics and heterospecifics might not be based on identical mechanisms. Indeed, it has been shown that *Drosophila* females manipulate and store sperm differently depending upon whether it is from conspecific versus heterospecific males (Manier et al. 2013a). Nonetheless, intraspecific sperm competition in *Drosophila melanogaster* is also well characterized at the genetic level (Begun et al. 2000, Findlay et al 2008), and it has been shown that genes mediating intraspecific sperm competition (Wolfner 1997, Neubaum and Wolfner 1999) also significantly affect the efficacy of conspecific sperm precedence (Castillo and Moyle 2014; Civetta and Finn 2014, and see Discussion). Pre- and postcopulatory remating success therefore offers a model for investigating mating traits in parallel for their role in both intraspecific and interspecific reproductive interactions.

Here we evaluated the potential link between these male reproductive traits in conspecific and heterospecific mating contexts, using a worldwide set of fifteen *Drosophila melanogaster* populations. To focus specifically on male performance traits, we examined remating success and offensive sperm competition in males from these different populations, all with a single female line and all against a standard conspecific and heterospecific first male genotype. We first evaluated evidence for variation among lines in pre- and postcopulatory male competitive traits against both conspecific and heterospecific males. With these data, we examined whether male success against conspecific first males was associated with success against heterospecific first males. By examining two different male performance traits, we were able to assess if either of these phenotypes meets the requirement for sexual selection to drive speciation—that these traits act in the same direction—and whether male reproductive success differences against conspecifics are alone sufficient to explain patterns of reproductive isolation. Finally, we also evaluated non-competitive male fecundity, to confirm that male sexual success was specifically due to male traits mediating competitive interactions. Based on our findings, as well as previous theoretical and empirical work, we identify several general conditions which might favor a positive relationship between sexual selection and reproductive isolation, and therefore enable a role for sexual selection in the evolution of strengthened reproductive barriers. Nonetheless, we conclude that even under such conditions, sexual selection is unlikely to be sufficient to drive reproductive isolation on its own.

## Methods

### Fly stocks and maintenance

All fly stocks were reared on standard cornmeal media prepared by the Bloomington Drosophila Stock Center (BDSC) at Indiana University, and were kept at room temperature (∼22C). Fourteen of our Drosophila melanogaster experimental male lines were drawn from the founder lines of the Drosophila Synthetic Population Resource (King et al. 2012; and provided by Stuart MacDonald) and were chosen as they represent a diverse sample of worldwide populations with genome sequence data for downstream studies (see Figure 1 for locations of origin). The fifteenth male line was the Austria w132 line, originally collected by Christian Schlotterer and donated to us by Kristi Montooth (University of Nebraska –Lincoln). This line was also used as the female genotype in all trials in this study. P1 male lines for intraspecific and interspecific competition trials were Green Fluorescent Protein (GFP) labeled *D. melanogaster* 32170 (obtained from the BDSC) and *D. simulans* 14021-0251.2663 (obtained from the University of California San Diego *Drosophila* species stock center), respectively. The same female line, and first male *D. melanogaster* and *D. simulans* GFP lines, were used in Castillo and Moyle’s (2014) analysis of the role of three known sperm competition loci in the expression of conspecific sperm precedence.

**Figure 1:**
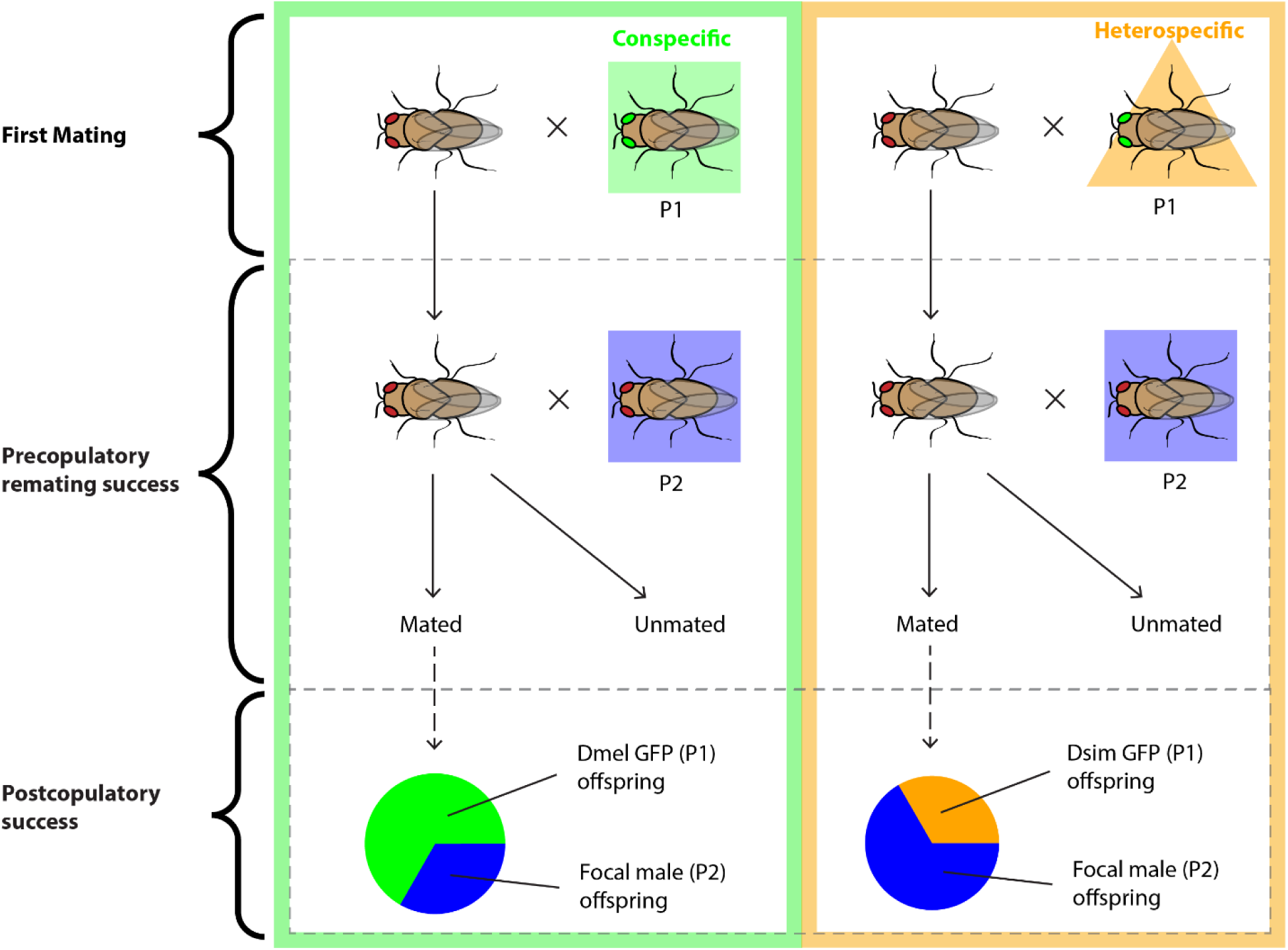
Schematic of experimental procedure. The experimental structure for assessing competitive success against conspecific (left) versus heterospecific (right) males is parallel, the only difference being the species identity of the first (P1) GFP-labelled male genotype. In each trial, a single female is first mated with a GFP-labelled first (P1) male before being paired with a second (P2) male drawn from one of the 15 target populations (Table 1). Adult offspring are scored for GFP ocelli after each second pairing, producing two datasets: precopulatory success (judged by presence/absence of wild-type (non-GFP) offspring) and, for pairings in which a second mating is confirmed, postcopulatory success (determined by proportion of offspring sired by P2 male). Both first male GFP genotypes and the female genotype - Austria w132 - are the same for all trials.

**Table 1:** Mean mating data for each male population. Precopulatory success is represented as proportion of total trials where P2 males succeeded in securing a remating, and postcopulatory success is shown as average proportion of total offspring that are sired by the P2 male. Noncompetitive fecundity is shown as average number of adult offspring produced in 24 hours following a single mating with each male line. Standard error is included for postcopulatory success and noncompetitive fecundity data.

|  |  | Precopulatory success |  | Postcopulatory success |  |  |
| --- | --- | --- | --- | --- | --- | --- |
| P2 male identity |  | Heterospecific first male | Conspecific first male | Heterospecific first male | Conspecific first male | Non-competitive fecundity |
| **A1** | Ohio | 0.727 | 0.625 | 0.942 ± 0.11 | 0.457 ± 0.03 | 23.8 ± 2.2 |
| **A2** | Colombia | 0.500 | 0.217 | 0.845 ± 0.13 | 0.436 ± 0.05 | 23.4 ± 1.7 |
| **A3** | Spain | 1.000 | 0.300 | 0.762 ± 0.05 | 0.095 ± 0.06 | 25.4 ± 1.0 |
| **A4** | Zimbabwe | 1.000 | 0.625 | 0.866 ± 0.06 | 0.354 ± 0.03 | 25.0 ± 1.2 |
| **A5** | Greece | 0.888 | 0.625 | 0.842 ± 0.09 | 0.222 ± 0.03 | 23.6 ± 0.7 |
| **A6** | Georgia | 0.667 | 0.556 | 0.867 ± 0.15 | 0.389 ± 0.06 | 22.6 ± 1.8 |
| **A7** | Taiwan | 1.000 | 0.714 | 0.930 ± 0.08 | 0.360 ± 0.02 | 26.0 ± 1.1 |
| **B1** | Bermuda | 0.714 | 0.625 | 0.893 ± 0.13 | 0.420 ± 0.03 | 19.8 ± 1.1 |
| **B2** | S. Africa | 0.875 | 0.375 | 0.885 ± 0.06 | 0.195 ± 0.03 | 20.2 ± 3.2 |
| **B3** | Israel | 0.667 | 0.500 | 0.935 ± 0.09 | 0.283 ± 0.03 | 21.8 ± 1.2 |
| **B4** | California | 1.000 | 0.857 | 0.935 ± 0.10 | 0.499 ± 0.02 | 23.8 ± 1.4 |
| **B5** | Hawaii | 0.889 | 0.667 | 0.955 ± 0.10 | 0.352 ± 0.01 | 25.2 ± 1.8 |
| **B6** | Peru | 0.900 | 0.556 | 0.959 ± 0.09 | 0.499 ± 0.01 | 25.2 ± 1.7 |
| **B7** | Malaysia | 0.900 | 0.333 | 0.933 ± 0.07 | 0.249 ± 0.02 | 24.8 ± 1.7 |
| **WT** | Austria | 1.000 | 0.857 | 0.893 ± 0.02 | 0.488 ± 0.02 | 22.8 ± 3.4 |

### Male intra- and interspecific performance assays

To assess differences in offensive competitive ability of males from our 15 target lines, our assay evaluated siring success of male lines with tester (Austria w132) *D. melanogaster* females following a first mating with either a GFP-marked male *D. melanogaster* (to assess intraspecific sexual competition) or *D. simulans* (to assess interspecific sexual interactions). The intra- and interspecific assays were designed so that the only difference between them was the identity of the first (P1) male, enabling us to directly compare the relative competitive success of each target male line against a common heterospecific and conspecific male tester genotype (Figure 1).

For first matings, virgin female Austria flies, GFP male *D. melanogaster* flies (for intraspecific trials), and GFP male *D. simulans* flies (for interspecific trials), were isolated 1 day prior to first mating. 4-6 virgin females and 4-6 males (either *D. melanogaster* or *D. simulans* P1 genotypes) were housed in a single vial and allowed to mate for 3 days. Females were then removed singly to individual blue-dyed food vials and allowed to oviposit for 24 hours, then checked for eggs to ensure at least one mating had occurred with a GFP-labeled first male. Females that did not oviposit were recorded as 'unmated' and removed from the remainder of the experiment (Figure 1). Females confirmed to have mated were then moved to a new vial to be singly paired with a virgin male from one of the 15 target male lines. The second pairing was maintained for 24 hours, after which the male was removed and the female moved to a new vial for 24 hours. Each female was then moved to a new vial every 24 hours for two additional days, for a total of 4 days of oviposition (3 of these after contact with second male). Progeny from all four vials were allowed to mature into adulthood and, upon eclosion, all adult individuals were gently anesthetized and viewed under a Leica M205FA stereo scope equipped with a UV light for visualizing GFP. The presence or absence of GFP in the ocelli of the eye was used as a marker of paternity; progeny with GFP ocelli must have been sired by the first mated (P1) GFP-labelled male, whereas progeny with wildtype ocelli are counted as progeny of the target male populations. For each timepoint per trial—the post-P1 blue vial and 3 post-remating vials—each progeny individual was scored for presence or absence of GFP in the ocelli. In instances where no wildtype offspring were observed in the 3 post-remating vials, we assumed that a successful remating event with the second male did not occur, and these were scored as ‘unremated’ (Figure 1). Both intra- and interspecific trials were repeated until at least 5 successful replicates (i.e. trials in which there was evidence that a second/remating event occurred) for each focal (second) male line were obtained.

### Estimating variation in first mating frequency

Because our experiment was designed so that post-conspecific and -heterospecific mating assessments were entirely parallel, the first pairing involved multiple males and females (4-6 each) for both conspecific (GFP *D. melanogaster* males) and heterospecific (GFP *D. simulans)* first matings. The latter was used to ensure that heterospecific matings occurred at a reasonable frequency, however a corollary effect is that females in conspecific first pairings might have experienced >1 copulations within the 3 day co-housing period. Because we did not directly observe matings, we used a secondary assay to estimate the number of conspecific males each female likely mated with during these first pairings, as follows: 5 virgin Austria females were paired with 5 conspecific males that were a mix of GFP males and males from a single focal line (either 2 GFP/3 focal, or 3 GFP/2 focal), and kept co-housed for 3 days as above. A total of 54 paired trials were run (half of 3 GFP/2 focal males, and half of 2 GFP/3 focal males). Within these 27 pairs of trials, focal males were drawn from four of our lines: Austria (10/27 pairs) and California (7/27 pairs), and Israel (5/27) and Spain (5/27). Following mating, females were allowed to oviposit for 24 hours in individual vials. Upon eclosion, progeny were assessed for presence versus absence of GFP in the occeli; each trial was then scored as ‘GFP’, ‘WT’ or ‘mixed’ matings based on the types of progeny found in the offspring. This procedure was not performed for heterospecific first matings, as the mating rate was low enough that females were assumed to be singly mated.

Assay results—the number of females that produced only monotypic versus mixed offspring— were used to estimate the likely number of copulations per female in these first mating trials, based on the general expectation that fewer mixed versus monotypic offspring broods is consistent with a lower frequency of females that have copulated more than once. We found that 16/54 trials produced ‘mixed’ offspring, all of which must have been the product of at least two copulations; the remaining females produced monotypic offspring (16/54 all GFP-labelled, and 22/54 all wild type). From these observations, and some simplifying assumptions (see Supplementary text), we infer that the majority of females mate either 1 or 2 times in first mating trial (for example, we estimate 48% are single matings), whereas the likely frequency of three copulations is low (Supplementary text).

### Quantifying male competitive success

We used success of second males in securing a remating as our data for offensive precopulatory success, for each male line. In particular, we tallied the number of second mating trials where there was at least one non-GFP offspring in subsequent vials—indicating a second mating with the target male—versus the number of trials that had only GFP offspring (unremated). The number of successes and failures were compared among male lines and between intra- and interspecific assays to determine if either second male line identity or identity of first male (conspecific vs. heterospecific) affected mean success in obtaining a remating (see analyses below).

We used the proportion of offspring sired after the second mating to estimate offensive postcopulatory (sperm competitive) success. For all trials in which a second (focal male) was successful at remating (i.e., for which there was at least one non-GFP labelled offspring in post-remating vials), the proportion of all adult flies that were wild-type versus GFP (tallied across the 3 post-remating vials) was calculated as an estimate of postcopulatory siring success, either in competition with conspecific (for intraspecific trials) or heterospecific (interspecific trials) males.

### Non-competitive male fecundity assay

To determine if our competitive male phenotypes were associated with non-competitive male fecundity, we assessed fecundity for each of the 15 target male lines when singly mated with the tester female line Austria w132. Virgin males and females were isolated and housed individually for 24 hours to reach maturity, prior to being aspirated into individual vials that contained one Austria female and a single male of a focal line. Each pair was then co-housed for 24 hours on standard media, after which the male was removed and the female allowed an additional 24 hours to oviposit before being removed. We assessed male fecundity at two stages—average number of eggs produced by an inseminated female and total adult offspring produced—in each assay, allowing us to also verify that adult progeny counts reflect successful fertilization events. Fecundity assays were performed for at least 5 replicates for each male line. We found that egg count and adult offspring count were quantitatively indistinguishable (i.e., with few exceptions, all eggs developed into adulthood), therefore non-competitive fecundity measures used in all subsequent analyses were based on adult offspring count (see Supplementary Text for egg data).

### Statistical analyses

#### Phenotypic variance for pre- and postcopulatory success

We used a χ^2^ test of independence to evaluate whether our 15 focal male lines differed in their mean precopulatory success at obtaining a second mating following a first mating with a tester conspecific male line (intraspecific trials), or following a first mating with a tester heterospecific male line (interspecific trials.) To evaluate whether male lines differed in postcopulatory offensive sperm competitive ability, we used analyses of variance (ANOVA) to assess whether identity of focal male genotype significantly affects the mean proportion of progeny sired by the second (P2) male. ANOVAs were performed separately for sperm competitive success following conspecific and heterospecific first matings, on a logit transformation of the proportion of offspring sired by the target male (i.e., not GFP labelled). To assess whether focal male lines differed in non-competitive fecundity, we used an ANOVA to assess whether focal male genotype significantly affected the mean number of adult offspring produced within 24 hours of a single non-competitive mating. For completeness, we also evaluated whether variation in either con- or heterospecific success at either mating stage was associated with global geographical patterns, and found that neither continent of origin, nor temperate versus tropical origin, significantly predicted competitive success in any of these cases (data not shown).

#### Relationship between male performance after intraspecific versus interspecific matings, and between pre- versus post-copulatory success

For both pre- and postcopulatory success traits we used Pearson’s correlation coefficient to assess the association between line means for reproductive phenotype following intraspecific versus interspecific first matings. For the male precopulatory success data, we assessed the correlation between proportion successful remating after conspecific trials versus after heterospecific trials, using line mean values. For postcopulatory success, we tested for a correlation between line mean siring success against conspecific versus heterospecific males. Within each class of interaction (intraspecific or interspecific) we also evaluated the correlation between pre- and postcopulatory success phenotypes using a Pearson’s correlation coefficient. Because assessing the correlation between mean phenotypes in these analyses does not take into account any within-line variance exhibited for postcopulatory phenotypes in our dataset, we also performed several bootstrap analyses to assess the effect of within- line (among-individual male) variance on the strength of these associations between male performance phenotypes (see Supplementary text).

#### Relationship between competitive and non-competitive male reproductive performance traits

To assess the relationship, if any, between non-competitive male performance and success in offensive reproductive interactions, we evaluated the correlation between male non-competitive fecundity and both pre- and postcopulatory success traits when competing against conspecifics. For each we computed a Pearson’s correlation coefficient between average fecundity of each male line (after single matings) against the logit transformation of proportion data for either pre- or postcopulatory success following a conspecific first mating. For completeness, we also examined the relationship between non-competitive fecundity and male pre- and postcopulatory success following a heterospecific first mating.

## Results

### Significant variation among male lines for reproductive success traits

We found significant phenotypic variance between focal male lines in their ability to secure a mating following conspecific (*χ^2^* = 24.784; *P* = 0.037) and following heterospecific (*χ^2^* = 23.804; *P* = 0.048) first matings (Figure 2). Following conspecific first matings, precopulatory success ranged from 0.217 to 0.857, with the majority of populations exhibiting a relatively high success rate (>0.6) but Colombia, Spain, South Africa, Israel, and Malaysia performing more poorly. The proportion of successful remating attempts following heterospecific males ranged from a perfect success rate (shared by 5 of the populations) down to 0.5 for the Colombia male line (Table 1).

**Figure 2:**
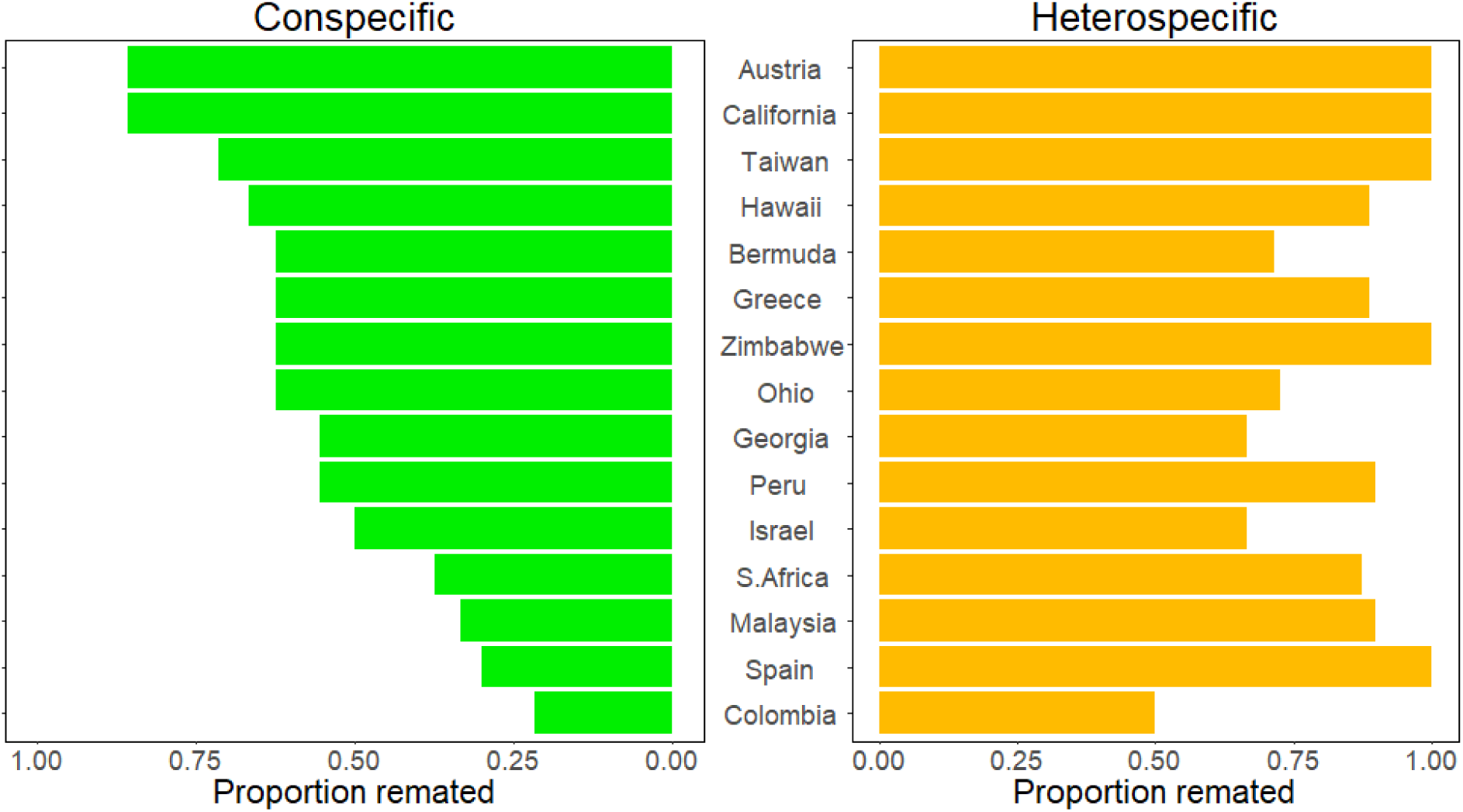
Precopulatory remating success following conspecific (left) and heterospecific (right) first matings. Precopulatory success is the proportion of remating trials that resulted in at least one non-GFP offspring, indicating the proportion of trials in which P2 males secured a remating. Male lines significantly vary in their precopulatory success following both conspecific (*χ^2^* = 24.784; *P* = 0.037) and heterospecific (*χ^2^* = 23.804; *P* = 0.048) first matings.

We also found significant phenotypic variation between focal male lines in their postcopulatory success as the second (offensive) male genotype, against both conspecific first males (*F*(14, 45.937); *P* = 0.022) and against heterospecific first males (*F*(14, 25.989); *P* = 0.007) (Figure 3). Interestingly, for postcopulatory success against conspecifics only, this was accompanied by a relatively high variance in performance among males within each line (Figure 3). Because this variation cannot be explained by male or female genotype effects, and the rearing environment of all flies was uniform, it might be due to variation among individual females in whether they had copulated once or twice prior to our remating assay (that is, within the initial 3-day pairing with conspecific tester males; see methods and supplement). Consistent with this, we see much less among-male within-line variation following heterospecific first matings, in which females are expected to only have mated once. Importantly, despite within-line male performance variation, we still detect significant line mean differences in pre- and postcopulatory success against the conspecific (GFP) tester male genotype.

**Figure 3:**
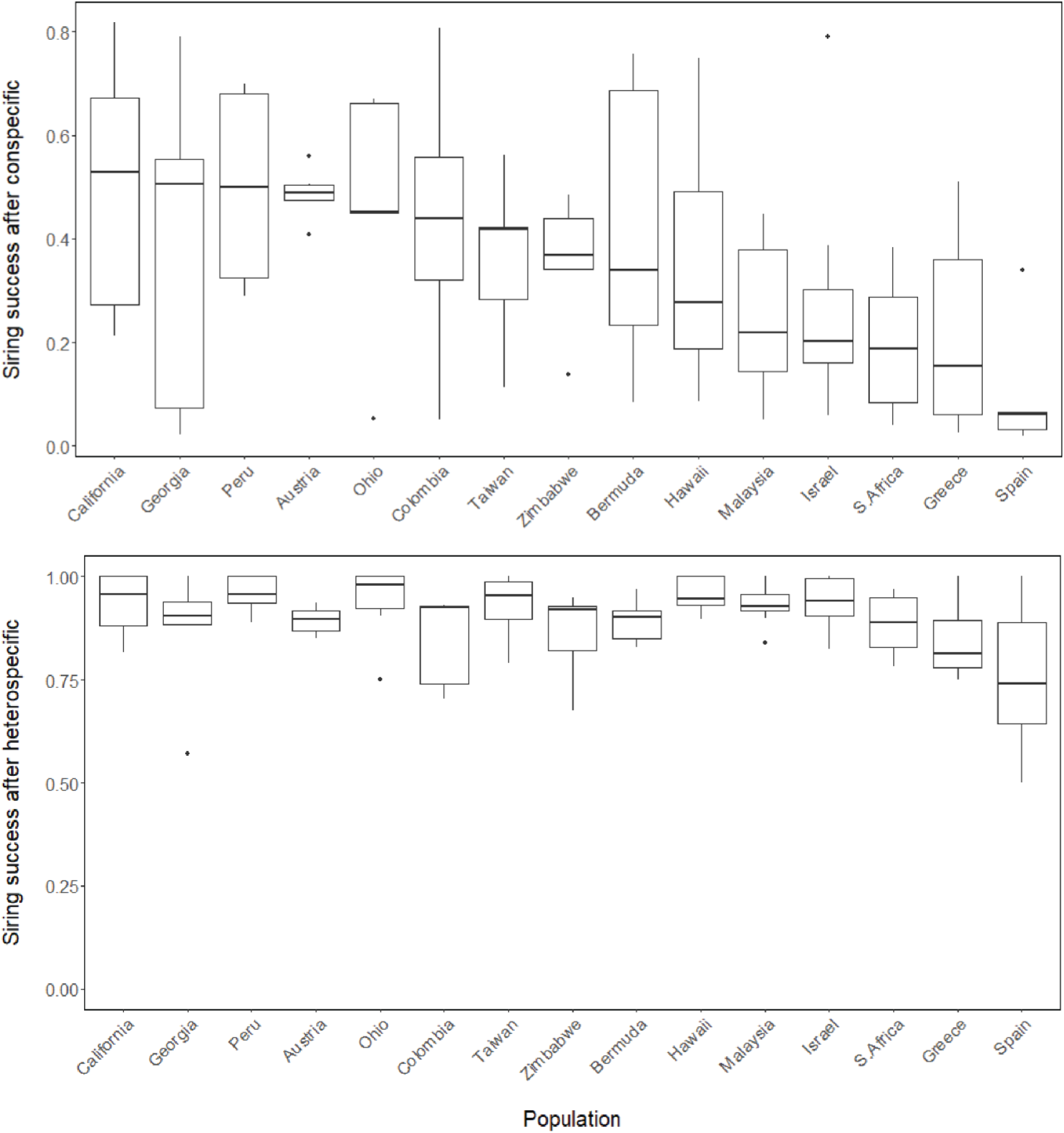
Postcopulatory siring success of second males from each focal line following intraspecific (top) and interspecfic (bottom) first matings. Boxes are mean proportion sired plus the 1^st^ and 3^rd^ quartiles; lines show standard errors. ANOVA on logit transformed proportions of offspring sired by the P2 male found significant male line effects for sperm competitive success after both conspecific (*F*(14, 25.989); *P* = 0.006877) and heterospecific (*F*(14, 45.937); *P* = 0.02192) first matings.

In contrast to competitive male reproductive phenotypes at both pre- and postcopulatory stages, we found no significant difference among male lines in non-competitive fecundity (*F*(14, 1.006); *P* = 0.459).

### Positive but modest relationship between male sexual performance against conspecific males and heterospecific males

To assess whether sexual selection and reproductive isolation could act in the same direction, we examined the relationship between success after conspecific versus heterospecific first matings for both pre- and postcopulatory traits. We found no significant correlation between precopulatory success after conspecific versus heterospecific first matings (r(13) = 0.450, *P* = 0.090)(Figure 4). In contrast, mean male line values of postcopulatory success against conspecific first males versus against heterospecific first males were significantly positively associated (r(13) = 0.552, *P* = 0.032)(Figure 5). This suggests that male lines that are on average better sperm competitors against conspecific males are also better able to displace heterospecific sperm.

**Figure 4:**
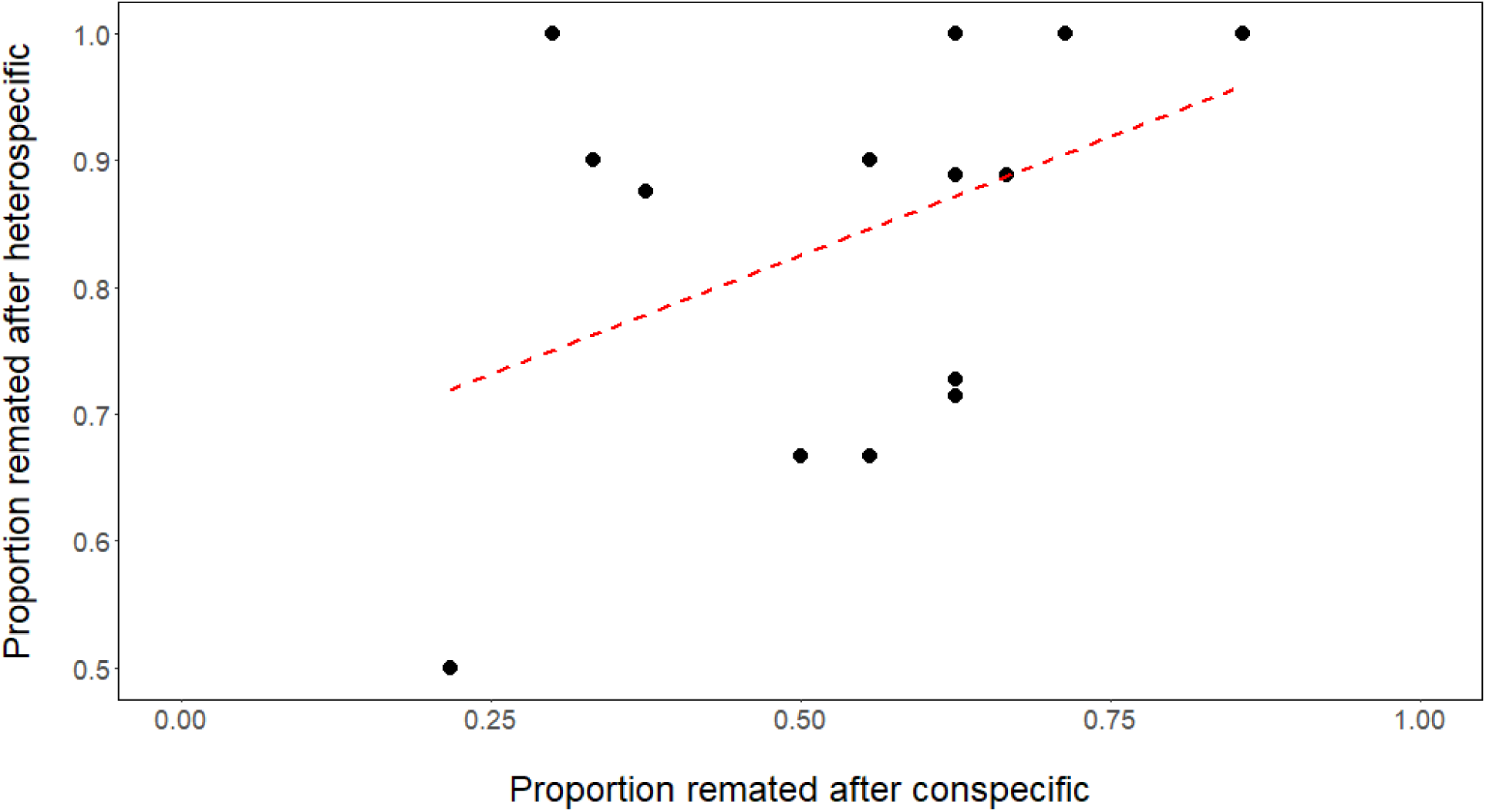
Mean precopulatory success of focal male lines after first matings with a conspecific (X-axis) versus heterospecific (Y-axis) first male is not significantly correlated (r(13) = 0.450, *P* = 0.090). Each point represents the proportion of second mating trials that were successful for a focal male line.

**Figure 5:**
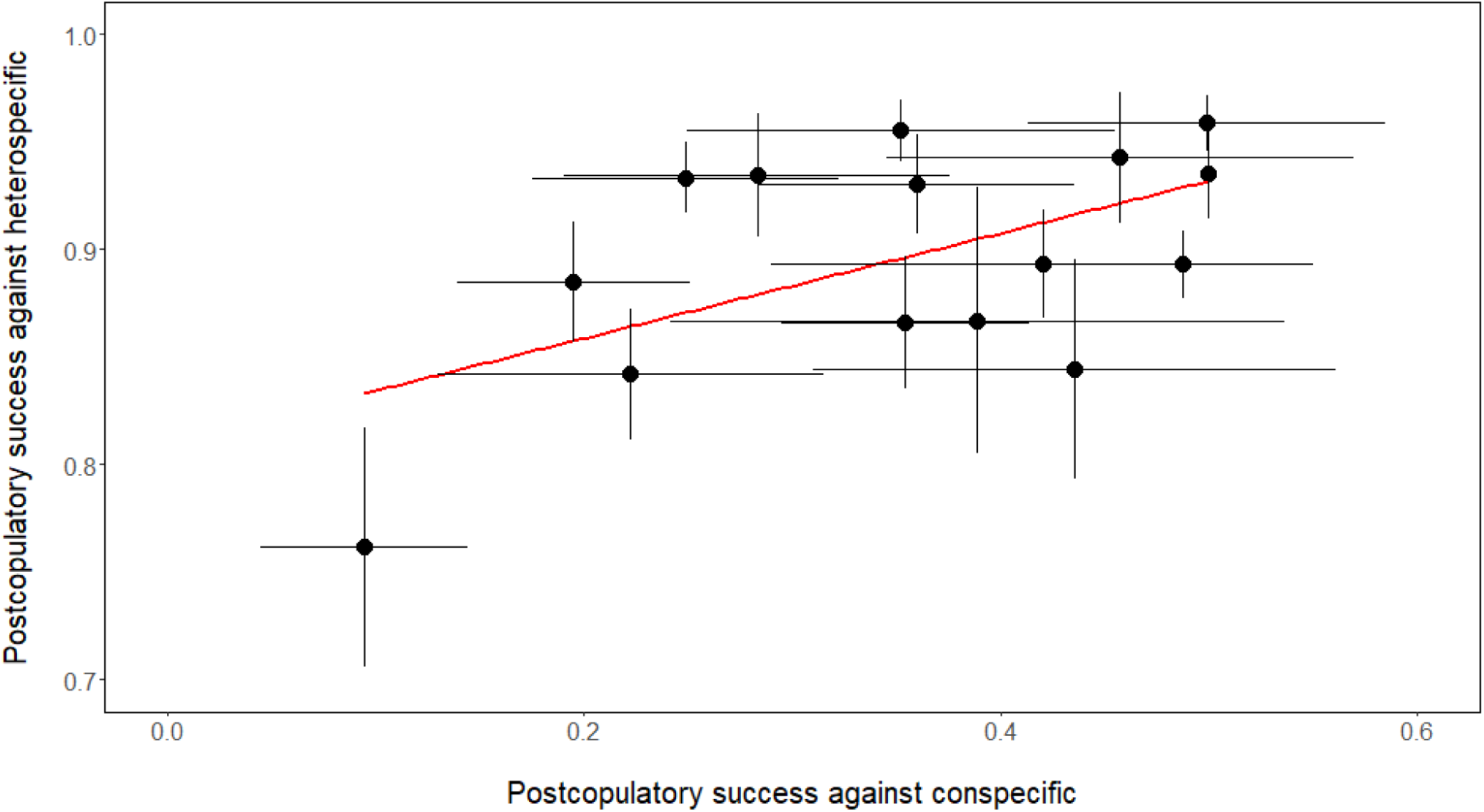
Mean postcopulatory competitive success of target male lines following first matings with conspecific (X-axis) or heterospecific (Y-axis) males is significantly correlated (r(13) = 0.552, *P* = 0.032). Points represent the average proportion of offspring sired by each male genotype, across all trials involving that genotype. The lines represent the standard errors for each population for the two phenotypes.

### Postcopulatory success is associated with precopulatory success after conspecific first matings

Because we assessed second male success in terms of both precopulatory and postcopulatory competitive ability, we could also evaluate whether these different sexual performance traits are associated with each other. For male performance after conspecific first matings, we found a significant correlation between remating success and sperm competitive ability (r(13) = 0.562, *P* = 0.029). In contrast, pre- and postcopulatory success following heterospecific matings were not associated (r(13) = 0.014, *P* = 0.959) (Figure 6).

**Figure 6:**
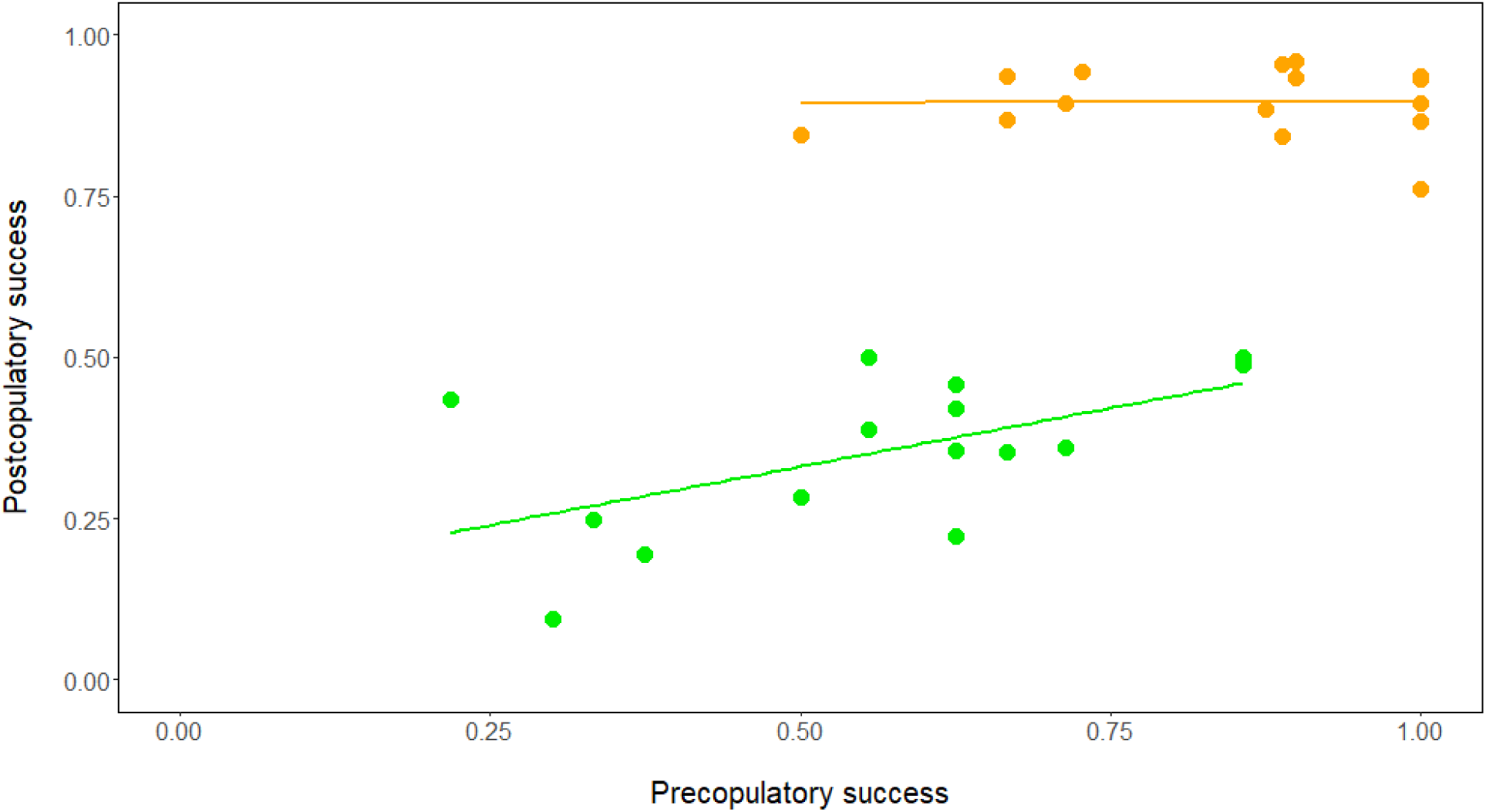
Associations between second male (P2) success at pre- and postcopulatory stages when competing against conspecific males (green) or heterospecific males (orange). For male performance against conspecifics, pre- and postcopulatory success is significantly correlated (r(13) = 0.562, *P* = 0.029); pre- and postcopulatory male performance following a heterospecific first mating is unassociated (r(13) = 0.014, *P* = 0.959).

### Differences in remating and sperm competitive success are not determined by non-competitive male fecundity

To better interpret the causes of variation in second male reproductive success, we evaluated whether male line variation in either precopulatory or postcopulatory success could be explained by differences in intrinsic, non-competitive male fecundity. We found no significant correlation between intrinsic fecundity and precopulatory remating success following conspecific (r(13) = 0.046, *P* = 0.870) or heterospecific (r(13) = 0.476, *P* = 0.073) first matings. We also found no correlation between noncompetitive fecundity and postcopulatory success against either conspecific (r(13) = -0.047, *P* = 0.868) or heterospecific (r(13) = 0.188, *P* = 0.503) first males.

## Discussion

Because of its effects on reproductive trait evolution, sexual selection is commonly invoked as a driver of speciation. However questions still remain about the mechanistic connection between conspecific and heterospecific sexual interactions, including whether sexual selection generally acts in the same or different directions as reproductive isolation. Here we examined phenotypic variation in, and the relationship between, pre- and postcopulatory male performance in *Drosophila melanogaster* when competing against a conspecific or heterospecific first male. Although we found no significant association between precopulatory success—securing a remating—after a conspecific versus a heterospecific first mating (Figure 4), average postcopulatory competitive success was significantly positively associated between the two contexts (Figure 5). Our findings speak both to whether sexual selection could be expected to act in concert with, or in opposition to, reproductive isolation, and whether selection within species is individually sufficient to drive reproductive isolation. They also raise the question of why the male performance traits examined here might differ in their degree of coupling between conspecific and heterospecific mating contexts, and suggest conditions under which these positive associations are expected to be more or less likely. Overall, our results indicate that sexual selection on some male reproductive traits can facilitate the evolution of isolating barriers, but may be insufficient to drive isolation on its own.

### Intraspecific competitive success has a positive but modest contribution to reproductive isolation

The specific connection between sexual selection and reproductive isolation determines whether sexual dynamics within species could shape the evolution of reproductive isolation between species, including facilitating or even constraining the emergence of isolating barriers. Our results indicate that, for the male performance traits we examined here, sexual selection does not appear to constrain reproductive isolation and could, for post-copulatory competitive traits, act to facilitate the expression of stronger reproductive isolation. Previous analyses have found mixed evidence for this association. While some theoretical work has suggested that sexual selection amplifies isolation under specific conditions (Gavrilets and Waxman 2002, Gavrilets 2000, Kondrashov and Shpak 1998, Higashi et al. 1999), other models indicate that sexual selection can oppose speciation, especially when strong female preferences reduce the variation in male mating traits that is required for reproductive isolation to evolve (Servedio 2012, Kirkpatrick and Nuismer 2004, Servedio and Burger 2014). Interestingly, most of the latter predictions have been generated specifically in the context of divergence with gene flow (i.e. in partial sympatry); it is unclear whether similar expectations hold broadly in allopatry, where there is no direct or indirect selection for increased reproductive isolation but also where gene flow— including the movement of strong female preference alleles between species (e.g. Servedio and Burger 2014) —does not oppose the build-up of isolation.

The few previous empirical assessments of relationships between sexually-selected traits and reproductive isolation have also produced variable findings. These differing outcomes mostly appear to depend on the specificity of female preferences for con- or heterospecific male traits in any given case. At least two studies have inferred that sexual selection works against reproductive isolation because an exaggerated male trait is preferred by both conspecific and heterospecific females in sympatry, thereby increasing the propensity for gene flow between species (*Girardinichthys*, Macias Garcia et al 2012; *O. pumilio*, Yang et al. 2016). In contrast, in both Laupala crickets (Mendelson and Shaw 2005, Shaw and Parsons 2002) and Tungara frogs (Boul et al. 2007), male signal traits that are strongly preferred within species are also used by females to discriminate against heterospecific males. While our analysis was not primarily designed to assess the pattern of female preferences for con- and heterospecific male traits, we found that the *D. melanogaster* female line we used generally chooses to remate with conspecific second males after a heterospecific first mating (i.e. more often than foregoing a second mating; Figure 4), and strongly prefers the sperm of second conspecific males after heterospecific first matings (i.e. siring success against *D. simulans* is well above 50% for all conspecific male lines; Figure 5), regardless of the significant variation among male lines in their relative success in each phenotype. Together, our observations and these prior studies suggest that strong female preferences for conspecific male traits could help to facilitate positive associations between sexual selection and reproductive isolation. Note, however, that female preferences need not always be required, especially in the specific context of sexual conflict over fertilization; for example, the sperm of *Caenorhabditis* lineages experiencing strong antagonistic selection on sperm competition traits have been shown to cause traumatic female sterility in heterospecifics (Ting et al. 2014), indicating that sexual antagonism acting within species can also produce isolation between species.

While our findings suggest that male sexual performance against conspecifics and heterospecifics can act in the same direction, at least for competitive fertilization traits, they also indicate a fairly modest association between sexually selected and reproductive isolating phenotypes. This observation implies that sexually selected intraspecific phenotypes can contribute to isolating barriers, but are likely insufficient to individually drive the reproductive isolation observed between species. Other studies that have investigated this general relationship—both theoretically and empirically—have made similar inferences (e.g. Bolnick and Kirkpatrick 2012). Using patterns of isolation observed in empirical examples, Bolnick and Kirkpatrick (2012) modelled the strength of intraspecific assortative mating that would be necessary to produce phenotypic differences sufficient to cause reproductive isolation, and showed that the magnitude of sexual selection fails to sufficiently explain the amount of reproductive isolation observed between lineages. Additional analytical theory and simulations suggest that male traits must be under both natural (divergent) selection and sexual selection in order to successfully contribute to isolation in sympatry (Kirkpatrick and Ravigne 2002; Servedio and Burger 2014; Servedio and Boughman 2017); sexual selection alone is insufficient to drive increases in isolation.

If sexual selection can be a positive but insufficient driver of isolation, it is important to examine the conditions under which sexual selection can be more effective (or less) at generating reproductive barriers. Interestingly, our data themselves suggest at least one other condition that could determine the strength of this association: the degree to which there are shared underlying mechanisms controlling competitive success against conspecifics versus heterospecifics. Below we discuss two lines of evidence that suggest that varying degrees of mechanistic overlap between these mating contexts likely explains both the modest association we observed for postcopulatory traits, and lack of relationship for precopulatory traits.

### The degree of mechanistic overlap influences the strength of association between sexually selected traits and reproductive barriers

Both of the sexual performance phenotypes we examined here are likely a product of multiple underlying mechanisms, that could vary in their degree of overlap between conspecific and heterospecific mating contexts and therefore in the expected strength of association between these contexts. Interestingly, evidence both from prior characterization of these pre- and postcopulatory performance phenotypes, and from direct genetic studies, suggests that the degree of mechanistic overlap could be greater for postcopulatory phenotypes. Prior analyses of the mechanics of postcopulatory sperm success in *Drosophila* indicate that it primarily acts on only three classes of trait: sperm traits (reviewed in Snook 2005), ejaculate traits (reviewed in Mautz et al. 2013) and female reproductive responses (including sperm transfer, sperm displacement, sperm ejection, and sperm selection for fertilization (Manier et al. 2013b, Manier et al. 2010, Miller and Pitnick 2002)). This circumscribed suite of traits increases the chance that superior male performance against conspecific and heterospecific males share some physiological mechanisms in common. Within *D. melanogaster*, for example, in addition to size and speed of sperm—traits that affect displacement success (e.g., Lüpold et al 2012)—intraspecific sperm success has been shown to vary in part due to female ejection timing (Lüpold et al 2013). Interspecifically, in *D. simulans* it has been shown that postcopulatory fertilization success of conspecific males over heterospecific males also depends on both male-mediated sperm displacement success and female-controlled mechanisms of sperm ejection, among other traits (Manier et al. 2013a). In addition, in *D. melanogaster* there is direct evidence that these postcopulatory mechanisms also share genes in common (Castillo and Moyle 2014). In assays using the same female line and the same first male genotypes as examined here, Castillo and Moyle (2014) showed that knockdown or null lines for two genes (*Acp36DE, CG9997)* with known roles in sperm competition within *D. melanogaster* were also significantly less competitive in offensive (second male) sperm competition with *D. simulans* first males. A third evaluated locus—*Sex Peptide*—is essential for effective sperm competition among conspecifics (Liu and Kubli 2013), but had no phenotypic effect on competitive performance against heterospecific first males (Castillo and Moyle 2014). Together, both direct phenotypic observations and genetic studies provide good evidence that there is some degree of overlap, although incomplete, in the mechanisms that mediate postcopulatory sperm success when competing against conspecific and heterospecific males.

In contrast, a much larger range of traits could mediate the precopulatory success of males in persuading *Drosophila* females to remate (reviewed Markow et al. 2005), including a complex set of non-independent behavioral cues (Welbergen et al. 1992) as well as several mating signals along at least two sensory axes - acoustic (reviewed Hoikkala 2005) and chemosensory (reviewed Chung and Carrol 2015) . Intraspecifically, *D. melanogaster* females are known to use both acoustic (e.g., pulse songs with more pulse (Talyn and Dowse 2003)) and chemical signals (e.g., presence and abundance of male pheromone 7-Tricosene; Billeter et al. 2009) to evaluate conspecific mating partners. How *D. melanogaster* females discriminate against heterospecific *D. simulans* males is less well understood (reviewed Sawamura and Tomaru 2002), however courtship song interpulse interval is known to affect interspecific courtship success (Kyriacou and Hall 1986). Given this broad variety of potentially important precopulatory traits, it is possible that mechanisms involved in male remating success after heterospecific versus conspecific first matings are not even on the same sensory axis. Moreover, the criteria for assessing both the benefit of remating and the suitability of second males, likely varies depending on the species identity of the first male. Following a heterospecific first male, females will almost certainly benefit from remating by compensating for an unambiguously low quality first mating (the “rescue effect”, Fricke et. al. 2006), whereas following a conspecific mating, the benefits of remating are more strongly dependent on whether the second male is a higher quality male (enabling the female to “trade up”, Byrne and Rice 2005). In the first instance, the criteria for identifying suitable second mates might be focused more on species-recognition traits, whereas the latter might assess more nuanced differences in male quality or persistence (although it remains unclear if mate recognition traits are generally distinct from species recognition traits; reviewed Servedio and Boughman 2017). Interestingly, our own observations following conspecific first matings suggest that females might indeed be making these remating decisions based on male quality; generally, male lines with better precopulatory remating success are also better sperm competitors against conspecifics following this mating (Figure 6). In contrast, the success of male lines in persuading females to remate following a heterospecific first mating is uncorrelated with their ability to displace heterospecific sperm (Figure 6). This discrepancy between these two associations is further evidence that overlap between mechanisms that underlie these traits is incomplete, as we might otherwise expect to see a similar pattern of success following a heterospecific mating as we do a conspecific if these mechanisms were identical. In either case, there are clear reasons that the degree of mechanistic overlap might be expected to differ systematically between traits involved in precopulatory and postcopulatory male success, when compared between conspecific and heterospecific first male contexts.

### Conditions most likely to connect sexual selection and reproductive isolation

As we've addressed above, our findings—in conjunction with previous theory and data—suggest several primary conditions where sexual selection within species is expected to have its largest role in reproductive isolation. In particular, when female preferences for intraspecific male traits is strong, these preferences are expected to also contribute to discrimination against heterospecific males. In addition, when a limited number of potential mechanisms underlie mating interactions, the likelihood that these mechanisms will be involved in both intra- and interspecific contexts is increased. The importance of pleiotropy between sexually selected traits and isolating barriers is a repeated and general inference from numerous models of sexual selection and speciation (Ritchie 2007), and the differences we observe between different mating traits (pre- versus post-copulatory) here might reflect the overall propensity for these different classes of traits to participate pleiotropically in both sexual selection and isolation.

Finally, apart from these conditions, there are other general factors that might influence the extent to which some sexually selected traits might be less reliable or consistent contributors to reproductive isolation. For example, while environmental conditions can play a role in postcopulatory mating traits (for example, by changing the number or quality of sperm produced, depending upon physiological condition; reviewed in Pitnick et al. 2009), there are reasons to expect that precopulatory traits (such as visual, auditory, and other sensory cues) are more directly influenced by external environmental factors. For example, elaborate courtship traits can make males more apparent to predators, a potentially important natural selective agent; similarly, the type and intensity of male signals can be shaped by the environment in which these signals must be transmitted (Boughman 2002). The extent to which sexual selection and natural selection interact to influence the evolution of isolating barriers is an area of substantial ongoing research (Boughman 2013, Safran et al. 2013), but clear examples from the sensory drive literature (Boughman 2001, Seehausen et al. 2008) and elsewhere (Hardwick et al. 2013) suggest that sexual selection on precopulatory mating traits might often take a back seat to direct natural selection on these same traits, a conclusion also supported by theory (Van Doorn et al. 2009, Servedio and Boughman 2017). The operation of additional (and potentially constraining) forces of natural selection on precopulatory traits is therefore another factor that might favor a stronger role for postcopulatory traits in mediating the connection between sexual selection and reproductive isolation.

Regardless, it is evident that in order for sexual selection to drive speciation, phenotypes associated with success in one context must also be successful in the other. Here we explicitly tested for a relationship between sexually selected (intraspecific), and reproductive isolating (interspecific) traits by comparing male success directly between the two contexts for two different male mating success traits. While we found little support that sexual selection would be sufficient to drive reproductive isolation on its own, postcopulatory competitive success appears to act in the same direction in both contexts. In conjunction with previous evidence that competitive fertilization within and between species also depends upon some shared genes (Castillo and Moyle 2014) and overlap in mechanisms of sperm displacement and cryptic female choice (e.g., Manier et al. 2013a) it is clear that sexual selection on this class of male competitive traits could contribute to elevated reproductive isolation between species, in the form of competitive heterospecific sperm displacement. In the absence of many empirical analyses of the direct association between traits important for success in both intra- and inter-specific interactions, it remains unclear if this observation applies more generally beyond *Drosophila melanogaster* remating traits. Nonetheless, our analysis shows that such direct tests are critical for assessing sexual selection’s relationship with reproductive isolation, including the relative role of sexual selection could play as a driving force in speciation.

## Acknowledgements

We thank Stuart MacDonald for providing DSPR founder strains for our use, Kristi Montooth for providing the Austria strain, and Dean Castillo both for help procuring fly strains and for advice on experimental design. Research was supported by Indiana University Department of Biology funding to LCM and JSD.

